# Faster inference of complex demographic models from large allele frequency spectra

**DOI:** 10.1101/2024.03.26.586844

**Authors:** Enes Dilber, Jonathan Terhorst

## Abstract

We present momi3, a new method for inferring complex demographic models using genetic variation data sampled from many populations. momi3 features many improvements over its predecessor momi2 (Kamm, Terhorst, Durbin, et al., 2020), including support for continuous migration, just-in-time compilation, and execution on GPUs; a standardized interface for specifying demographic models; and a novel importance sampling strategy that enables it to efficiently analyze data from a large number of samples. Together, these improvements lead to speedups of as much as 1000× over existing state-of-the-art methods such as ∂*a*∂*i*, moments, and momi2. We illustrate the usefulness of our method by revisiting a model of archaic admixture using a large, recent dataset containing hundreds of human genomes from many populations.

## 1 Introduction

It is now widely appreciated that patterns of population genetic variation can be used to recover detailed information about the timing and magnitude of population size changes, migration, and admixture events, a pursuit known as *demographicinference* (Marchi, Schlichta, and Excoffier, 2021). Recently, the amount of data available for performing demographic inference has experienced substantial growth, along three different dimensions. First, the number of samples collected from individual populations continues to grow at a fast rate, due to the ever-decreasing cost of sequencing genomes. Second, the number of populations that have been sampled has itself also sharply increased (The 1000 Genomes Project Consortium, 2015; Mallick et al., 2016), owing in part to growing recognition of the scientific and ethical importance of sampling underrepresented groups. Finally, as a result of impressive technical achievements in ancient DNA extraction and analysis (Prüfer et al., 2014; Green et al., 2010; Meyer et al., 2012), we now have access to the genomes of individuals who lived at many different points in time as well.

New data sources should, in principle, enable us to answer increasingly precise and subtle questions about evolutionary history. However, in order to realize such a goal, lingering technical obstacles must be surmounted. Evaluating the likelihood of raw sequence data under completely realistic biological and evolutionary models is infeasible, even for small sample sizes (N. Li and Stephens, 2003; Sheehan, Harris, and Yun S Song, 2013; Rasmussen et al., 2014; Terhorst, Kamm, and Yun S Song, 2017). Therefore, approximations or data reduction strategies have to be employed in order to scale demographic inference methods up to the size of modern data sets.

In this article, we focus on one such strategy, which is to first summarize genetic variation data into a low-dimensional tensor, and then find an evolutionary model which would have generated a similar statistic. The summary statistic we focus on is known as the (joint) site frequency spectrum (JSFS). With *n* exchangeable samples from a panmictic population, the SFS is an (*n* − 1)-dimensional vector that counts of the number of singletons, doubletons, tripletons, etc. that were observed. More generally, if there are *n* samples from each of *p* populations (or “demes”), the JSFS is a tensor of dimension *O*(*n*^*p*^).

Exponential scaling in the number of populations means that state-of-the-art SFS-based demographic inference methods like ∂*a*∂*i* (Gutenkunst et al., 2009), momi2 (Kamm, Terhorst, Durbin, et al., 2020), or moments (Jouganous et al., 2017) are limited to analyzing, in a practical amount of time, perhaps several dozen individuals sampled from about five demes. Unfortunately, this falls well short of the quantity data that is now available. Wohns et al. (2022), for example, recently published a unified dataset containing over 3,600 human genomes from 215 populations—far in excess of what any tool we are aware of can analyze. Researchers face the unpleasant choice of either throwing away information to get an answer quickly, or waiting a large amount of time to perform a complete analysis.

To address this shortcoming, we have developed a new, scaleable method called momi3 for inferring complicated demographic models using modern data sets. It features many improvements over its predecessor momi2. First, the internal code has been completely rewritten using neural network libraries designed for performing large-scale machine learning on graphs (Bradbury et al., 2018). This brings several benefits, including end-to-end automatic differentiability, just-in-time compilation, and the ability to run on an accelerator (GPU/TPU) with zero additional effort. Second, support for modeling and inferring continuous migration has been added, thus lifting one of the main technical restrictions of momi2. Third, momi3 features a novel approximation technique, based on importance sampling, which can drastically reduce computational requirements for analyzing complex, multi-population demographic models, with a minimal impact on accuracy. Finally, we have eliminated large amounts of custom API in favor of standardized, community-developed libraries: `tskit` for storing and accessing genotypes; `msprime` (Baumdicker et al., 2022) for conducting simulations; and `demes` (Gower et al., 2022) for demographic model specification and visualization. Taken together, these advances enable momi3 to fit models containing hundreds of sampled individuals spread across many populations, in a computationally efficient and intuitive manner.

## 2 Background

Formally, we want to solve the following estimation problem: given allele count data 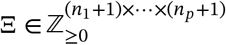 from *p* different populations, find a demographic model Θ existing in some model class ℳthat maximizes the likelihood of the data:

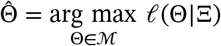

Under further assumptions (namely, linkage equilibrum and infinite sites) needed to render the problem tractable, it can be shown (Bhaskar, Wang, and Y. S. Song, 2015) that the maximum likelihood problem is equivalent to minimizing the Kullback-Leibler divergence between the observed frequency spectrum, and its expected value 𝔼_Θ_Ξ (when both are normalized to form discrete probability distributions):

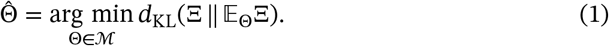

### 2.1 Prior work

Given Ξ and 𝔼_Θ_Ξ, computing the KL divergence is trivial; the difficulty of the problem lies entirely in evaluating the expectation. At least three different strategies have been proposed. First, simulation-based methods (Excoffier and Foll, 2011; Excoffier, Dupanloup, et al., 2013; Excoffier, Marchi, et al., 2021) use Monte Carlo estimation to numerically approximate 𝔼_Θ_Ξ. These have the advantage of being very flexible and requiring fewer simplifying assumptions; hence, the class of models (the ℳ in equation 1) is larger, and includes any process that can be forward-simulated. However, as Monte Carlo methods, they suffer from the curse of dimensionality, and encounter trouble when the number of populations *p* is large. In particular, in large samples the JSFS tensor Ξ is extremely sparse (see examples below), so, unless many iterations are used, the Monte Carlo estimate of 𝔼_Θ_Ξ has a different sparsity pattern, and the objective function (1) is not even defined.

As an alternative to simulation-based approaches, several exact methods have also been proposed. There are basically two approaches. Diffusion-based methods work by solving a system of differential equations which characterize the distribution of a biallelic variant over time in the infinite population limit. Two well-known implementations of this technique are the programs ∂*a*∂*i* (GGutenkunst et al., 2009; Gutenkunst, 2021) and moments (Jouganous et al., 2017). The second class of methods uses coalescent theory to compute 𝔼_Θ_Ξ. This line of work originates in a seminal paper by Griffiths and Simon Tavaré (1998), who showed how to compute the expected frequency spectrum for a single population whose historical size varied over time. Chen (2012) and Chen (2013) later extended this result to multiple, non-admixing populations, as did Kamm, Terhorst, and Yun S Song (2017), using different ideas which led to a much faster algorithm. Most recently, Kamm, Terhorst, Durbin, et al. (2020) further generalized the method to the case of admixed populations in the program momi2.

The coalescent and diffusion approaches possess different, and somewhat complementary, advantages and disadvantages. The running time of the latter is independent of sample size, and the diffusion can more easily incorporate non-neutrality and migration. However, convergence issues can arise when integrating the differential equations, and they are limited in the number of populations they can analyze—the original version of ∂*a*∂*i* could handle up to three; this was later increased to five using GPUs (Gutenkunst, 2021). moments removes some of these limitations by solving an alternative system of differential equations, but is also limited to no more than five populations, and its running time scales with sample size. Another important difference between the approaches emerges in large data sets: diffusion methods essentially solve for the entire joint frequency spectrum, even though most entries are not observed in real data, resulting in much wasted computation. For their part, coalescent-based methods are able to compute individual SFS entries, but have a computational requirement that grows with the number of samples, and cannot easily accommodate selection or continuous migration.

### 2.2 New contributions

momi3 is a significantly updated version of our earlier method momi2 which is oriented towards analyzing modern, large-scale population genetic datasets in a timely fashion. In addition to being much faster than existing methods, momi3 is able to combine the advantages of both the diffusion and coalescent approaches to demographic inference using the JSFS. The next few sections explain how we are able to achieve this.

#### 2.2.1 Summary of momi2

First, we need to review some salient technical details about how its predecessor works. momi2 decomposes an arbitrary demographic model containing splits, mergers, pulse admixture events, and population size changes, into a data structure called an *event tree*. Probabilistic message-passing (Wainwright and Jordan, 2008) is then performed on this tree in order to compute the likelihood of a given configuration of ancestral and derived alleles sampled at the tips of the demographic model. A complete technical description of the momi2 algorithm can be found in Kamm, Terhorst, Durbin, et al. (2020). We follow their notation (see Table 1 of that paper for a guide) in the sequel to streamline the exposition and minimize confusion.

**Table 1:**
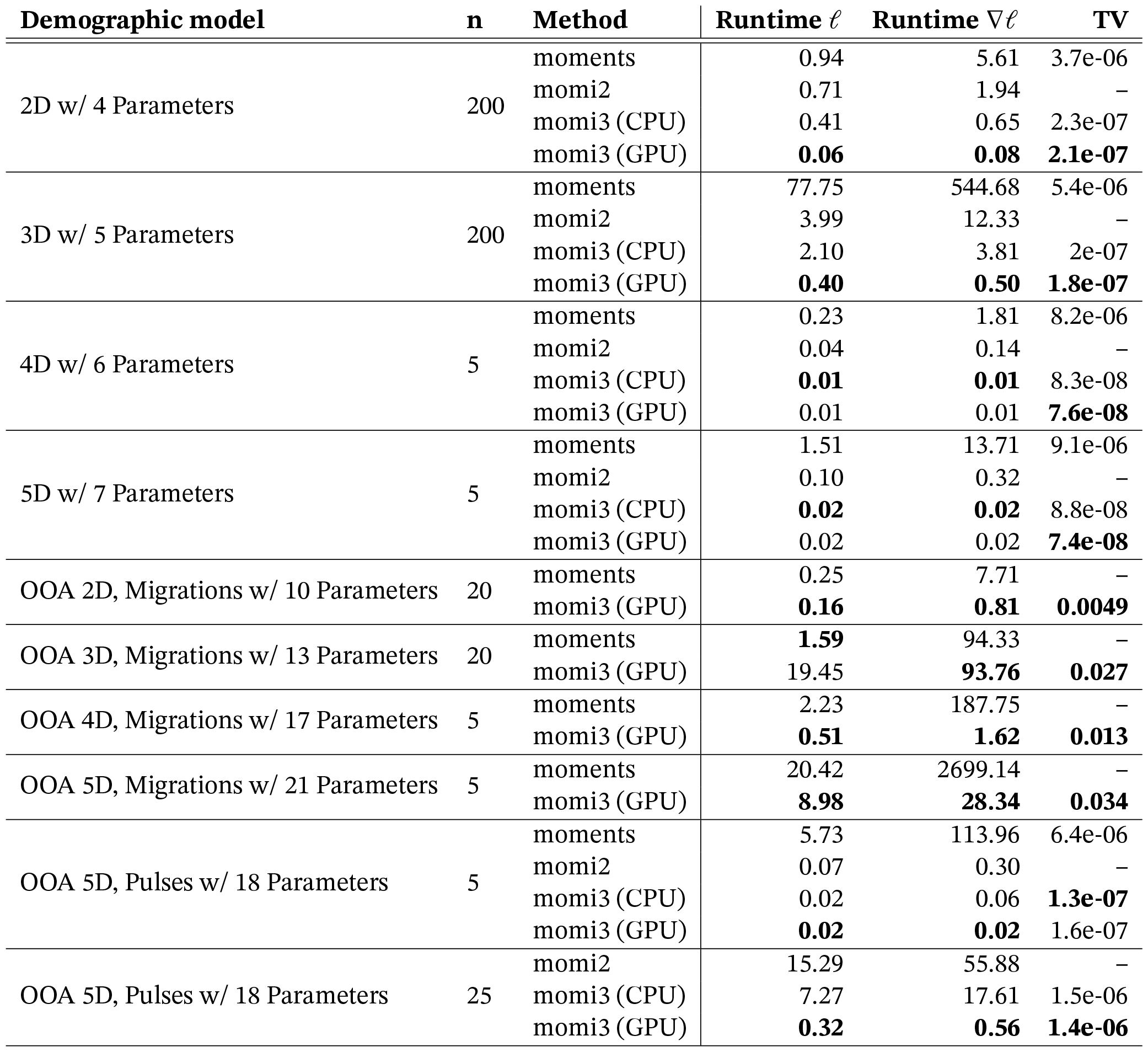
Computation time of the likelihood (𝓁) and the gradient (∇𝓁) along with Total Variation (TV) from the reference method for several demographic models. 15 CPUs have been used in each CPU test and a single GPU for each GPU test. Runtime ∇𝓁 is the total computation time of derivative of all variables and the likelihood.

To compute the expected frequency spectrum, momi2 requires two quantities. First, there is the partial likelihood tensor 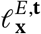, which, for a given node *E* in the event tree, gives probability of all of the data in the clade subtended by *E*, conditional on the state ***x*** of the ancestral populations at *E*. The second is the “truncated SFS”, *f*_*v*_(*k*), for a given population *v*, which is defined by the relationship

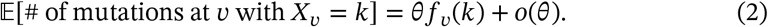

In words, *f*_*v*_(*k*) is the expected number of mutations, under an infinite sites model, that have *k* ancestral copies in *v* at the earliest (closest to the present) time at which *v* exists in the model. momi2 works by solving a dynamic program, denoted DP(𝓁^1^, …, 𝓁^*p*^) (c.f. Kamm, Terhorst, Durbin, et al., 2020, Algorithm 2), in terms of 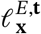 and *f*_*v*_(*k*) for each population *v* and event *E* in the event tree representation of the demography.

#### 2.2.2 Continuous migration

momi2 allowed for specifying (subject to memory and computation constraints) arbitrary numbers of “pulse” admixture events between populations. In this type of event, each surviving ancestral lineage present in a particular source population migrates independently with probability *p* to a specified destination population. However, mathematical difficulty prevented us from incorporating the continuous flow of migrants between populations. In momi3, we have added support for continuous migration, thus lifting one of the main technical restrictions of momi2.

To achieve this, we build on recent work by Jouganous et al. (2017) who used a moment-based approach to speed up the diffusion calculations necessary for computing the expected SFS forwards in time. The fundamental quantity in moments is expected number of segregating sites which are observed **i** ∈ (*n*_1_ + 1) × … × (*n*_*p*_ + 1) times in a sample of size **n** = (*n*_1_, …, *n*_*p*_) (denoted Φ_**n**_(**i**) in their paper.) As shown by Jouganous et al., as this tensor evolves forwards in time, it satisfies the system of ordinary differential equations

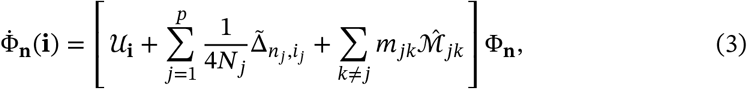

where the multilinear operators 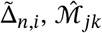, and 𝒰_*i*_ are given by equation (A2), equation (A4) (using the jacknife approximation described in Appendix D), and Appendix B, respectively, of Jouganous et al. (2017).

As already noted by Gravel and co-authors, the moments quantity Φ_**n**_ is quite related to the partial likelihoods tracked by momi2; they are, in a sense, “dual” to each other. To see this, observe that the “drift operator” 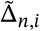 in (3) is precisely the *transpose* of the Moran rate matrix utilized by momi2 (compare equations (A2) of Jouganous et al. with the definition of the rate matrix *Q* = (*q*_*ij*_) immediately preceding equation (19) of Kamm, Terhorst, and Yun S Song, 2017). This is so because time runs oppositely in the two algorithms: momi2 transitions partial likelihood tensors *backwards* in time by right-multiplication with *e*^**Q**^, where **Q** = *Q*_1_ ⊕ … ⊕ *Q*_*p*_ is the Kronecker sum of Moran rate matrices corresponding to each population *i* = 1, …, *p*. Similarly, moments transitions the expectation tensor Φ_**n**_ *forwards* in time by solving the system (3), which reduces (up to scaling factors) to Φ_**n**_ = **Q**^T^Φ_**n**_ in the case of constant population size and no mutation or migration.

For a given event tree node *E*, let *K*_*E*_ be the set of populations that exist at *E*, as in Kamm, Terhorst, Durbin, et al. (2020). To lift the partial likelihood of a set *M* ⊂ *K*_*E*_ of populations experiencing continuous migration τ generations into the past, we solve the *adjoint* of (3) subject to the initial condition 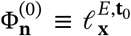, where **t**_0_ = **t** − τ∑_*v*∈*M*_ **e** _*v*_, to arrive at 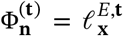 In practice this simply amounts to taking the transpose of each of the operators shown above and solving the resulting ODE system. To solve for the adjoint when all populations have constant size, we use a matrix-free method for computing the action of the matrix exponential (Al-Mohy and Higham, 2011) on the initial tensor 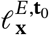, which allows us to exploit the sparsity and/or Kronecker product structure of the operators 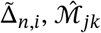 and 𝒰_*i*_ via fast matrix-vector products. If one or more populations experiences growth, then the *N*_*j*_ in terms (3) are no longer constant, so we use a differential equation solver as in Jouganous et al.

The other quantity we require is the truncated frequency spectrum, denoted *f*_*v*_(*k*) above in the case of a single population *v*. To generalize this to the setting of continuous migration, we must compute the truncated frequency spectrum for multiple populations exchanging migrants—a new quantity not considered in momi2. To find it, we simply observe that the expectation on the right hand side of equation (2) is exactly the same as the quantity Φ_**n**_ tracked by moments. Hence, differentiating the solution of the original system of ODEs obtained by moments with respect to the mutation rate parameter yields the multipopulation analog of the truncated frequency spectrum. Note that, in contrast to the lifting operation, here we solve the primal system forwards in time. The appropriate initial condition is thus 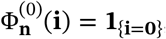, which ensures that only branch length arising from *de novo* mutations in the populations in question are considered in the expectation. The following proposition formalizes these arguments.

##### Proposition 1

*Let* 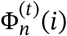 (*i*) *be the solution to the system* (3) *under a symmetric finite-genome mutation model (i*.*e. μ* = ν = θ *in the operator* 𝒰_*i*_), *with the initial condition is* 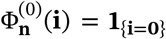. *The truncated frequency spectrum up to time* τ *for these populations equals*

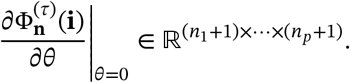

To compute the derivative displayed above, we can use either automatic differentiation (see below), or solve a related system of ODE’s using the forward sensitivity method (e.g., Bartlett, 2008).

#### 2.2.3 Genealogical importance sampling

Mathematically, momi2 computes the likelihood of a genetic variant by integrating over all possible genealogies on which the variant could have arisen. Often, this integral includes portions of the genealogical state space that are extremely unlikely, and contribute negligibly to the end result. For example, in a demography that features many pulse admixtures, momi2 expends computation to consider the scenario where *every* lineage migrates from destination to source population at *every* admixture event. Similarly, in a population which has experienced recent exponential growth, i.e. most human populations, momi2 allows for the possibility that *none* of the present-day samples have reached common ancestors, even thousands of generations back into the past.

Under plausible evolutionary models, these genealogies have vanishing likelihood, and could be dropped from the computation with little loss of accuracy. To improve the efficiency of momi3, we implemented a form of importance sampling which automatically prunes unlikely genealogies from the integration, thereby lessening both the memory and computational requirements of analyzing large samples. The basic observation is that the dimensionality of the Moran process which underlies our approach can be greatly reduced conditional on the event that only a small number of lineages are ancestral to the sample at a given time. To make this rigorous, let us introduce some additional notation. For a population *v* ∈ {1, …, 𝒟 } in the demography, let 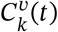 denote the event that the population immediately ancestral to *v* has no more that *k* lineages ancestral at time τ. Then we have the following result:

##### Proposition 2

*Let* 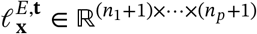 *be a partial likelihood tensor, where p* = |*K*_*E*_|, *and let v* ∈ *K*_*E*_ *be a population included in the partial likelihood tensor. Then conditional on the event* 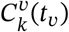 *that v has no more than k surviving ancestral lineages at time t*_*v*_, *the output of* DP(𝓁^1^, …, 𝓁^*p*^) *is unchanged if we make the substitution*

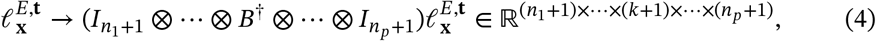

*where* 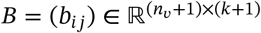 *is amatrix whose* (*i, j*)*-th entry is the probability of sampling without replacement j* ∈ {0, …, *k*} *derived alleles in the population immediately ancestral to v out of a total of i* ∈ {0, …, *n*_*v*_} *derived alleles segregating in v, and B*^†^ *denotes the Moore-Penrose pseudoinverse*.

*Proof*. The correctness of the modified algorithm follows directly from Lemmas 4 and 7 of Kamm, Terhorst, Durbin, et al. (2020): although those results are stated in terms of the partial likelihood of the leaf nodes (**X**_Leaves(*E*)_ in their notation), by the Markov structure of the multipopulation coalescent, they also apply if we consider the partial likelihoods at internal nodes of the event tree. Hence, conditional on 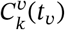, the population immediately ancestral to *v* has at most *k* descendants, so by Lemma 7, its partial likelihood is independent of allele counts of the remaining lineages {*k* + 1, …, *n*_*v*_}. Finally, the combinatorial relationship between 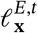 the partial likelihood of the population immediately ancestral to *v* is given by Lemma 4 (see also the discussion immediately preceding the statement of the lemma).

Using the proposition, we can substantially reduce the computational requirements of momi3 by heuristically searching for populations in the event tree where 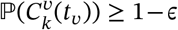 for some *k* ≪ *n*_*v*_ and ϵ ≪ 1.

In our current implementation, we focused on two types of events that are likely to result in a large reduction of the number of lineages that must be tracked. The first is transitioning (“lifting”) a partial likelihood backwards for large amounts of coalescent time. A lifting operation (cf. Lemma 1 of Kamm, Terhorst, Durbin, et al., 2020) involves transitioning a partial likelihood from a more recent to more ancient time using the Moran genealogical process. Suppose we are lifting a particular population *v*, with effective size *N*_*v*_(*t*) and sample size *n*_*v*_, from time *t* to time *t* + Δ*t*. Define

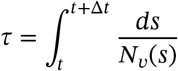

to be the amount of scaled coalescent time that elapses during this operation. Finally, let 𝒜_n_(*t*) denote the coalescent ancestral process associated to this population, i.e. a pure death process with 𝒜 _*n*_(0) = *n* almost certainly, and death rate *μ*_*i*_ = *i*(*i* − 1)/2 when the process is in state *i*. To find *k* and ϵ as above, we simulate^1^ the coalescent ancestral process a large number of times, and find the 1 − ϵ quantile of the random variable 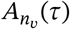. The correctness of this procedure follows directly from formula (2.1) of Griffiths and Simon Tavaré (1998).

A similar approach can be used to lessen the computational requirements for models involving admixture. Going backwards in time, a pulse admixture event causes a population *v* (say) to split into two parent populations of sizes *n*_*v*_ each, because it is possible that as few as none, or as many as all, of the lineages descended from either parent population. Let *p* be the expected fraction of lineages in *v* that came from parent 1, with the remainder coming from parent 2. The total number of lineages in *v* that came from parent 1 is distributed Binomial(*n*_*v*_, *p*). Hence, by considering the tail probabilities of the binomial CDF (which are exponentially small for *k* ≫ *n*_*v*_*p*), we can again find *k* such that 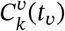 occurs with probability close to 1.

It is important to note a few caveats and tradeoffs. In order for the approximation error to be low, we need ϵ to be sufficiently small. Indeed, if ϵ = 0 then 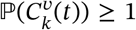 only when *k* = *n*_*v*_, so no approximation error is introduced and no computational savings are realized. Increasing ϵ trades accuracy for computation time. We explore this in more detail in simulations below. Additionally, at least some time needs to elapse before populations interact with each other in order for the Proposition to have application. In particular, it cannot currently be used to simplify computations for large populations that continuously exchange migrants all the way up to the present.

#### 2.2.4 Implementation improvements

Our implementation of momi2 used a combination of Python and hand-tuned C code. Although this approach yielded state-of-the-art performance, it had several disadvantages, such as its reliance on now outdated automatic differentiation libraries, and difficulty with porting the method to run on graphics processing unit (GPU) hardware.

momi3 has been rewritten from the ground up using modern machine learning libraries designed for performing large-scale tensor computations on graphs. We selected the probabilistic programming language Jax (Bradbury et al., 2018) due to its combination of speed, ease-of-use, and support for the Numpy API already utilized by momi2. This brings several benefits. First, momi3 natively supports automatic differentiation and just-in-time (JIT) compilation, leading to speedups as high as 1000-fold, as shown below. Second, the implementation seamlessly runs on CPU, GPU, as well as Google tensor processing (TPU) hardware, requiring no more than a command-line switch to select between the different backends. A third, more diffuse benefit, is that the speed and stability of momi3 will continue to improve over time, due to ongoing development of Jax by Google engineers and open source contributors. Indeed, we have already observed this; while we were preparing this article, support for sparse tensor computations in Jax improved considerably, leading to performance gains in momi3 with no additional effort on our part.

Readers should keep in mind that this approach requires a one-time compilation step, which can require several minutes for complex demographic models. This step must be repeated any time the structure of the event tree changes. Typically, the cost of compilation will be amortized over many of evaluations of the likelihood function and/or its gradient, however this may not always be true. One scenario where compilation time might become excessive is when searching over many topologically distinct demographic models, since recompilation would need to happen for each new topology. We discuss the cost of compilation in the simulation examples below.

#### 2.2.5 User interface improvements

Finally, we have made momi3 easier to use by relying on community-maintained libraries and standards wherever possible. First and foremost, we have eliminated momi2’s custom API for demographic model specification in favor of `demes` (Gower et al., 2022). Users can perform inference simply by providing demography model in `demes` format, along with the data, and model estimates can also be returned as `demes`-formatted demographies. Similarly, momi3 can directly interface with `msprime` (Baumdicker et al., 2022) for demographic model simulation and bootstrapping, and can directly read succinct tree sequence data (Kelleher, Etheridge, and McVean, 2016) using tskit.

momi3 is implemented using pure Python 3, with minimal dependencies and no additional compilation required. Source code and installation instructions are available at github.com/jthlab/momi3.

## 3 Results

### 3.1 Performance of momi3 vs. existing methods

In this section, we compare the performance of momi3 versus our earlier software momi2, as well as moments, which is the state-of-the-art diffusion-based method for this problem. We measured both running time and accuracy. Benchmarks were carried out using nine different demographies. The models denoted *xD*, for *x* ∈ {2, 3, 4, 5}, are demographies containing *x* isolated populations with no migration or admixture, and constant population size. The OOA models are all realistic Out-of-Africa models and have as parameters those estimated previously by Ragsdale and Gravel (2019). Only for the model with pulse migration, we added two pulse migrations to the demography tree for testing purposes. Visualizations of each of the models are shown in Figures S1 and S2. Summaries of all of the parameters for each model are in Tables S1 and S2. All of the simulations used a simulated chromosome of length 100Mbp.

#### 3.1.1 Running time

First we studied the time needed for each method to evaluate the log-likelihood of a particular model, and/or its gradient. For the CPU-based benchmarks, we ran all methods on unloaded cluster nodes containing Intel Xeon Gold 6254 CPUs. Since all the CPU-based methods can use multiple cores, we allocated 15 cores to each method when performing the evaluation (with one additional core occupied by the benchmarking code itself.) For the GPU runs involving momi3, we used a single NVIDIA Tesla V100 GPU.

Table 1 contains the running times for each method under these settings. The runtime of likelihood calculation can be seen at the column “Runtime 𝓁” and likelihood+gradient calculation in “Runtime ∇𝓁”. We calculated the gradients for each parameter in the given model. We used the built-in automatic differentiation capabilities of momi2 and momi3, while for moments, we relied on numerical differentiation as implemented in their software package.

First, we consider the performance improvement of momi3 relative to its predecessor momi2. For constant size models (demographies that begin with *xD*), momi3 running on a single GPU is more than 10 times faster in likelihood calculation, and up to 20 times faster in gradient calculation. For models featuring pulse migrations, the improvements are even more pronounced, e.g. a 30-fold improvement OOA-5D-Pulses demography. Less dramatic performance improvements on the order of 2-5× over momi2 were also observed even when we ran momi3 on CPU only, due to the efficiency gains of JIT compilation.

Next we turn to comparison with moments. In general, the simulation results show that large efficiency gains are again possible using momi3, however the improvements are not uniform. The largest differences were observed when comparing the time needed to evaluate the gradient of the 3D model with sample sizes *n* = 200, where momi3 was about 1000-fold faster than moments. This is because diffusion-based methods compute all entries of the joint frequency spectrum, whereas momi and other coalescent-based approaches compute them entrywise, so they can exploit sparsity in the observed SFS. For instance, in the 3D model with *n* = 200 samples per deme, moments computes all 201^3^ ≈ eight million SFS entries, even though only about 10^4^ distinct SFS entries are observed in a chromosome-length simulation.

Figure 1 shows how running time increases for each method on this demography as both sample size and the number of SFS entries increase. In Figures S3 and S4, we repeated this analysis for larger 4*D* and 5*D* models, but we were not able to include moments, since with e.g. 5 demes it would need to compute ≈ 3.5 × 10^8^ SFS entries. Overall, for simple demographies that do not have have admixture or migration, momi3 running on GPU can be expected to perform exceptionally well compared to moments.

**Figure 1.**
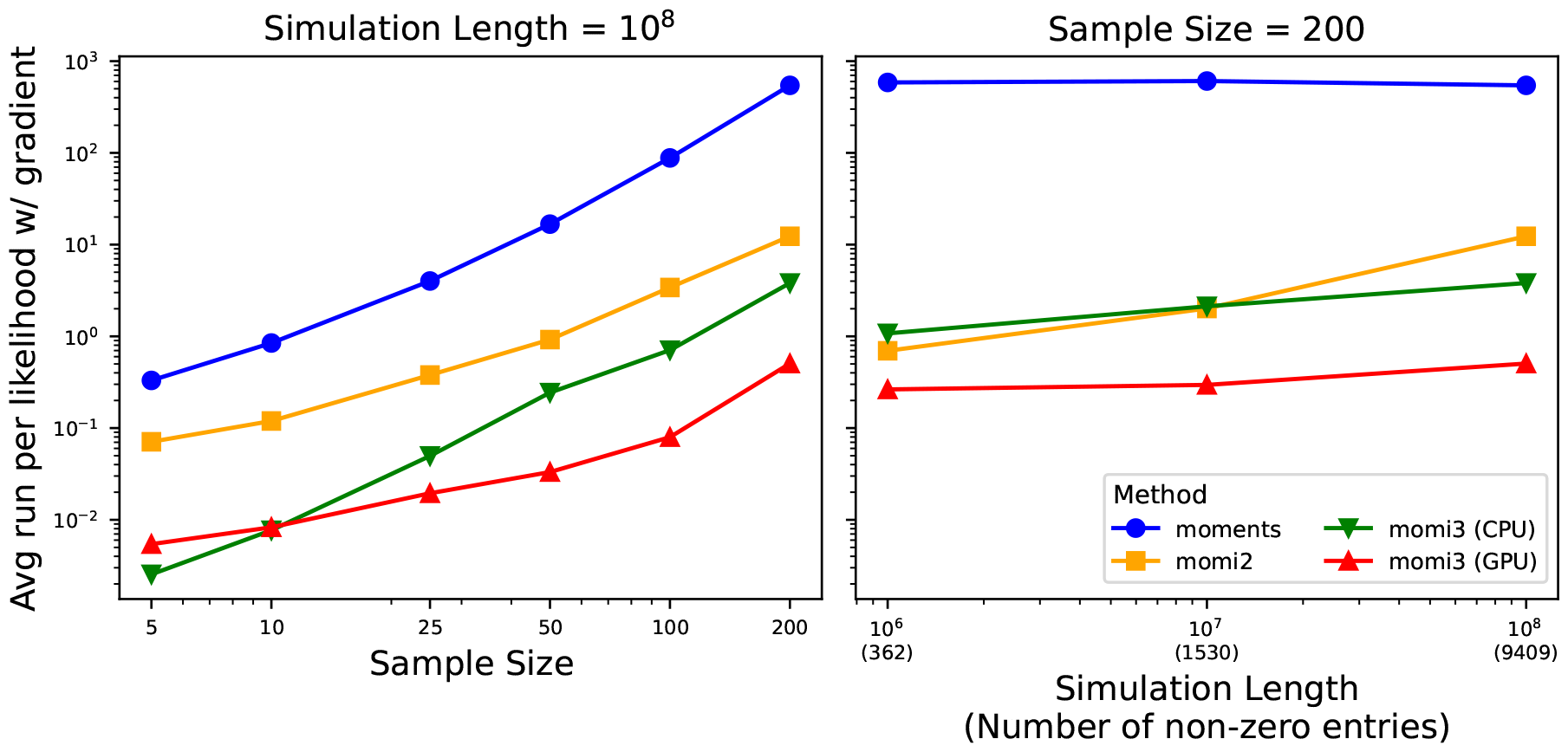
Runtime of a likelihood with gradient for the demography *3D w/ 5 Parameters* (See Figure S1). The figure shows how each method scale by the sample size (left) and simulation length (right).

For models involving admixture or migration, we found less dramatic improvements, but in most cases momi3 was the fastest method. The biggest differences were observed when computing the likelihood+gradient, where for complex models (e.g., OOA 5D with either pulse or continuous migrations), momi3 was hundreds of times faster than moments; this was mainly attributable to the inherent speed advantage of automatic (momi3) over numerical (moments) differentiation. However, there were also examples such as OOA 3D where moments was faster than momi3 even though the latter was running on a GPU. For continuous migration models, most of the execution time for both methods is spent solving differential equations via sparse matrix multiplication. Support for sparse matrix routines in Jax is currently experimental, and we expect that the performance of momi3 will continue to improve as Jax matures.

#### 3.1.2 Numerical accuracy

The last column of Table 1 shows the numerical accuracy of each of the methods. For the accuracy metric, we used total variation (TV) distance between the computed and true expected frequency spectrum (viewed as a probability distribution over allele configurations). Since TV is between 0 and 1, this can be interpreted as percent difference between the two distributions. To compute the true frequency spectrum, we must of course rely on another computational method, chosen to minimize numerical error. For demographies that do not involve continuous migration, we used the values returned by momi2 as the reference, since the method is exact for that class of models. For demographies involving continuous migration, we followed moments.

The results indicate that momi3 typically had the lowest numerical error for demographies involving pulse migrations. In fact, the difference between momi3 and momi2 was usually about *O*(10^−8^), which is the limit of floating point accuracy when using 32-bit floating point numbers, the default mode for momi3. For larger, more complex demographies the error was sometimes as high as *O*(10^−6^), which is still minuscule and should not affect most inference problems. These results are expected because momi2 and momi3 are algorithmically equivalent for this class of demographies, even though their implementations differ.

For demographies that have continuous migration, momi3 had error of about 1% − 3%, depending on the complexity of the demography. This was about an order of magnitude larger than the difference between moments and ∂*a*∂*i*. We expected greater numerical differences to emerge in this setting because the method of calculation is fundamentally different; momi3 needs to compute a derivative (see Proposition 1), which will tend to introduce numerical inaccuracy compared to methods that do not. Potential users of momi3 should keep this in mind, and decide whether somewhat lower accuracy is compensated for greater speed and scaleability.

#### 3.1.3 Compilation time

Finally, we benchmarked the compilation time of momi3 for each of the demographies listed above. As noted above, JIT compilation is unique to momi3 and a source of large efficiency gains. However, the compilation can require several minutes depending on the complexity of the demography. Enabling gradient mode also further increases compilation time. Table 2 shows compilation time (in seconds) for each demography, combination of likelihood or likelihood+gradient, and backend (CPU or GPU). For models that do not involve continuous migration, compilation times are quite reasonable; even the largest models we analyzed, containing hundreds of samples and many demes, compiled in under a minute. Compilation times tended to be slightly higher for GPU than CPU, and significantly higher for models that feature continuous migration. In the latter case, a lot of additional code is added including an ODE solver and many instances of (experimental, unoptimized) sparse matrix multiplication. Efforts to reduce compilation times for Jax models (for example, by caching compilation results and modularizing subroutines) are currently underway, so we expect that compilation times will improve in the future.

**Table 2:** Compilation time of momi3.

| Demographic model | n | $\ell$-CPU | $\ell$-GPU | $\nabla\ell$-CPU | $\nabla\ell$-GPU |
| --- | --- | --- | --- | --- | --- |
| 2D w/ 4 Parameters | 5 | 0.32 | 0.56 | 3.24 | 4.51 |
|  | 10 | 0.37 | 0.57 | 3.69 | 4.31 |
|  | 25 | 0.36 | 0.65 | 3.57 | 4.56 |
|  | 50 | 0.44 | 0.70 | 4.06 | 4.84 |
|  | 100 | 0.55 | 0.70 | 4.05 | 4.82 |
|  | 200 | 0.89 | 0.78 | 4.71 | 5.87 |
| 3D w/ 5 Parameters | 5 | 0.44 | 0.76 | 3.79 | 5.16 |
|  | 10 | 0.50 | 0.79 | 3.97 | 5.00 |
|  | 25 | 0.53 | 0.91 | 4.11 | 5.31 |
|  | 50 | 0.78 | 0.92 | 4.73 | 5.80 |
|  | 100 | 1.00 | 0.99 | 5.07 | 6.87 |
|  | 200 | 2.58 | 1.41 | 8.50 | 13.13 |
| 4D w/ 6 Parameters | 5 | 0.59 | 0.96 | 4.26 | 5.63 |
|  | 10 | 0.68 | 1.01 | 4.75 | 5.72 |
|  | 25 | 0.77 | 1.18 | 4.70 | 6.59 |
|  | 50 | 1.20 | 1.16 | 5.78 | 7.05 |
| 5D w/ 7 Parameters | 5 | 0.76 | 1.20 | 4.87 | 6.40 |
|  | 10 | 0.85 | 1.22 | 5.08 | 6.42 |
|  | 25 | 1.03 | 1.41 | 5.57 | 7.34 |
|  | 50 | 1.72 | 1.47 | 7.07 | 9.01 |
| OOA 2D, Migrations w/ 10 Parameters | 5 | – | 9.83 | – | 62.67 |
|  | 10 | – | 10.24 | – | 66.51 |
|  | 15 | – | 10.57 | – | 66.31 |
|  | 20 | – | 11.82 | – | 72.70 |
| OOA 3D, Migrations w/ 13 Parameters | 5 | – | 16.47 | – | 97.17 |
|  | 10 | – | 18.45 | – | 107.65 |
|  | 15 | – | 24.08 | – | 127.60 |
|  | 20 | – | 43.08 | – | 232.78 |
| OOA 4D, Migrations w/ 17 Parameters | 5 | – | 40.09 | – | 222.25 |
| OOA 5D, Migrations w/ 21 Parameters | 5 | – | 78.36 | – | 396.78 |
| OOA 5D, Pulses w/ 18 Parameters | 3 | 4.57 | 6.02 | 15.84 | 20.93 |
|  | 5 | 4.86 | 6.15 | 16.62 | 21.16 |
|  | 10 | 5.49 | 6.23 | 18.47 | 22.87 |
|  | 15 | 6.51 | 8.51 | 21.03 | 28.49 |
|  | 25 | 12.56 | 8.27 | 37.73 | 40.71 |

### 3.2 Real data applications

Next, we turn to some applications of momi3 on real data. As shown in the preceding section, momi3 basically produces the same output as existing methods, but is faster and able to analyze larger samples. The papers describing those methods already performed extensive real data analysis, and we do not expect to find anything new by reproducing their results. We therefore focused primarily on analyzing real data problems that are out of the reach of existing methods.

#### 3.2.1 Performance of the approximate method

First, we show how the approximation strategy outlined in Section 2.2.3 can be used to greatly reduce the computational burden of analyzing complex demographies using many samples. For this section we analyzed an out-of-Africa model with Neanderthal admixture (Figure 4). We fit this model using all of the available samples in the dataset published by Wohns et al. (2022): 107 Yoruba, 28 French, and 1 Vindija Neanderthal genomes, for a total (haploid) sample size of 272. For simplicity, we used data from chromosome 20, and focused on inferring three parameters: τ_5_, the time of the out-of-Africa event; η_1_, the population size of Ancient Modern Humans; and π_0_, the admixture proportion of Neanderthals into the OOA population.

Due to the large number of pulse admixtures, computations involving the exact model are slow. The first row of Table 3 (“No Sampler”) shows that, after a 26-second compilation step, each evaluation of the likelihood and its gradient takes roughly 3.1 seconds. Even for this simple, 3-parameter model, maximizing the likelihood requires several hundred gradient steps, so fitting time is several minutes overall. To quantify uncertainty by bootstrapping, we would need to repeat this step hundreds of times, requiring a large amount of computational resources.

**Table 3:**
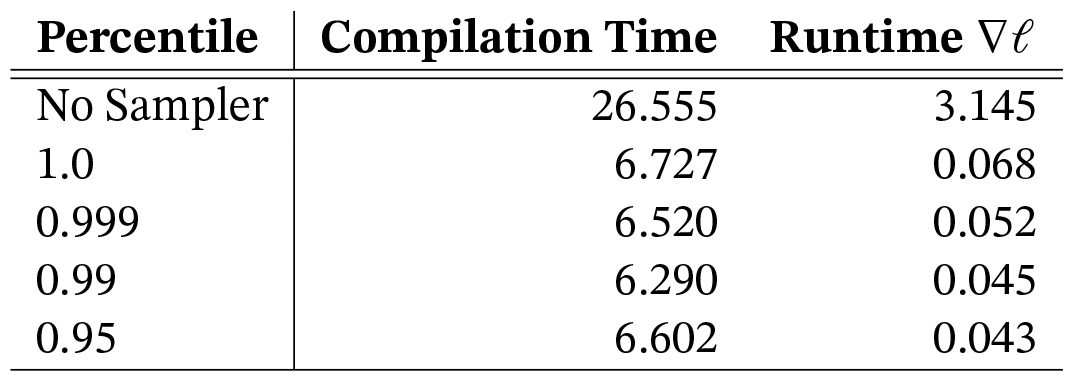
Running times using genealogical importance sampling.

This motivated us to study how Proposition 2 could be applied to improve performance. For each tolerance level ϵ ∈ {0, 0.001, .01, .05}, we performed Monte Carlo simulations to estimate the 1 − ϵ quantile for the number of surviving ancestral lineages at each node in the event tree representation of this demography, and then used the built-in downsampling option of momi3 to perform approximate inference. This strategy produced a speedup of roughly 50-75× over the exact method (Table 3 rows 2–5^2^), with larger ϵ leading to greater speedups. We then assessed the bias and variance of the estimated 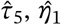 and 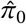. Here bias is defined as the difference between the maximum likelihood estimate using the exact model, versus each of the downsampled models, across a large number of bootstrap resamples from the observed frequency spectrum. Figure 2 shows the distribution of relative bias for each setting of ϵ (violin plots of each individual estimator are shown in Figure S6). With 1 −ϵ ≥ .999, estimates using the approximate models are virtually indistinguishable from the exact model across all parameters. However, larger values of ϵ introduced more error. For η_1_, using 1 − ϵ ≤ .99 produced a bias of about 1%, equivalent to about 150 units of *N*_*e*_, and similar results were observed for estimating the admixture fraction π_0_. Interestingly, the divergence time estimate τ_5_ seemed to be relatively less affected by downsampling, with mean bias of only about 0.1%.

**Figure 2.**
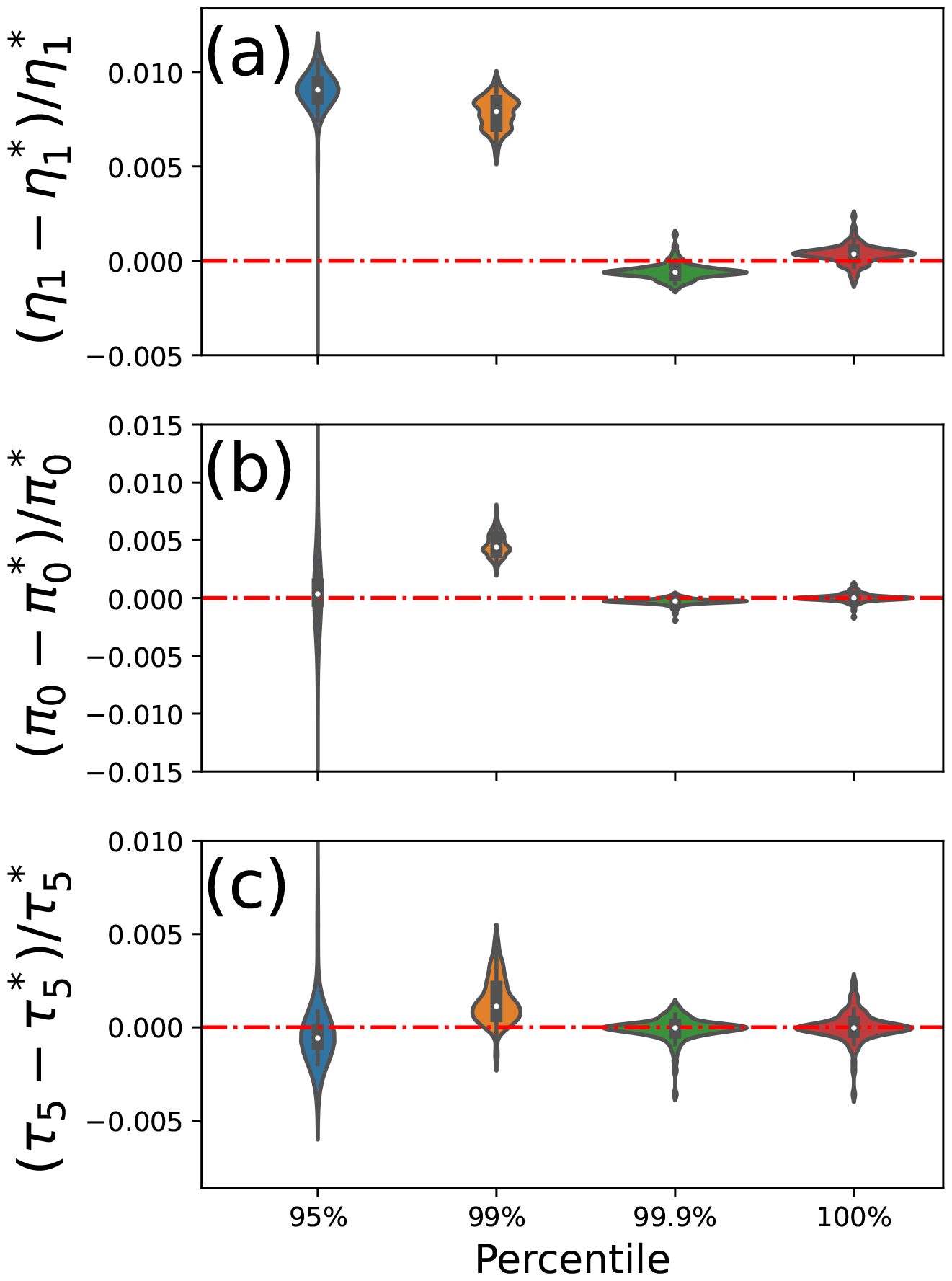
Distribution of the bias on the estimates of Figure 4 model in violin plots. Distributions are obtained by bootstrap samples. Star variable represents the No Sampler estimates for the given bootstrap sampler. Closer to red line is better.

In summary, using large values of ϵ (particularly ϵ = 0.05) introduces additional bias and variability, and is probably best avoided except as a last resort. However, setting ϵ = .001 (or even ϵ = 0) resulted in a substantial efficiency gain, with minimal impact on the parameter estimates.

#### 3.2.2 Inference of archaic admixture using many genomes

Encouraged by these results, we turned to a more complex demographic model of archaic admixture. As is well-known, non-African individuals carry about 1-4% Neanderthal DNA due to past interbreeding between humans and Neanderthals. There is also evidence of admixture from Denisovans into modern humans. However, the precise nature and timing of these events is uncertain and the subject of ongoing investigations (Sankararaman et al., 2014; Vernot and Akey, 2015; Dannemann and Racimo, 2018) . In particular, it is not clear if these admixture events are better modeled a sequence of discrete pulses, or perhaps as low-level continuous gene flow over a period of time (Browning et al., 2018; Villanea and Schraiber, 2019; Durvasula and Sankararaman, 2020).

To study this question we used the demographic model proposed by Reich et al. (2010) and fit it to all the avaliable data in Wohns et al. (2022) dataset (all sample sizes are diploid): 107 Yoruban samples, 28 French, 15 Papuan, as well a Vindija Neanderthal, and a Denisovan individual. Figure 3 shows a visualization of this demography. We could not use either ∂*a*∂*i* or moments to analyze this model, since it contains more than 5 populations. Nor could we employ momi3 directly; attempting to do so exhausted all the available memory on our GPU. Thus, we turned to importance sampling, as in the preceding section, in order to make progress.

**Figure 3.**
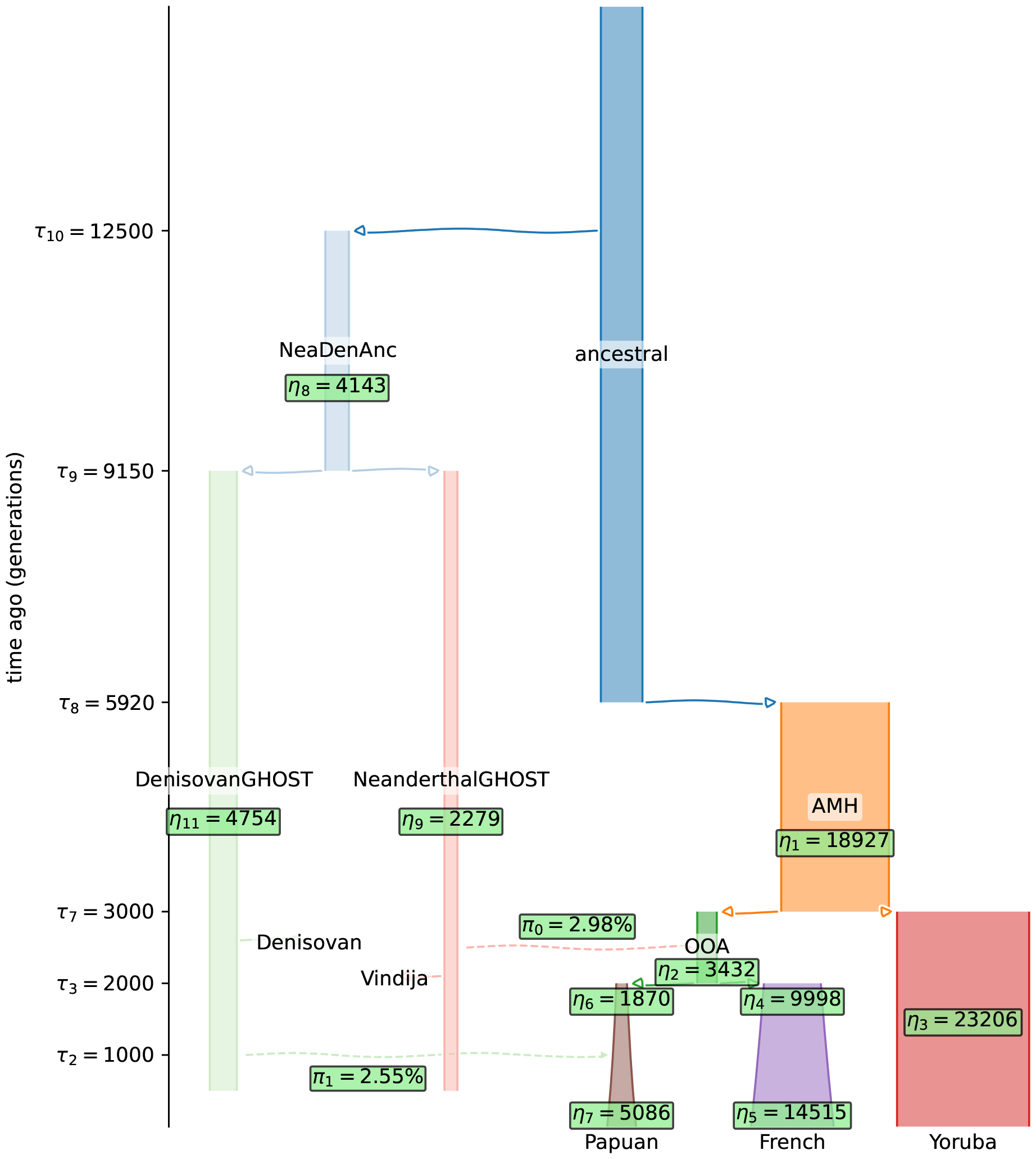
Out-of-Africa model with two extinct hominids. Parameter estimates are shown in green.

**Figure 4.**
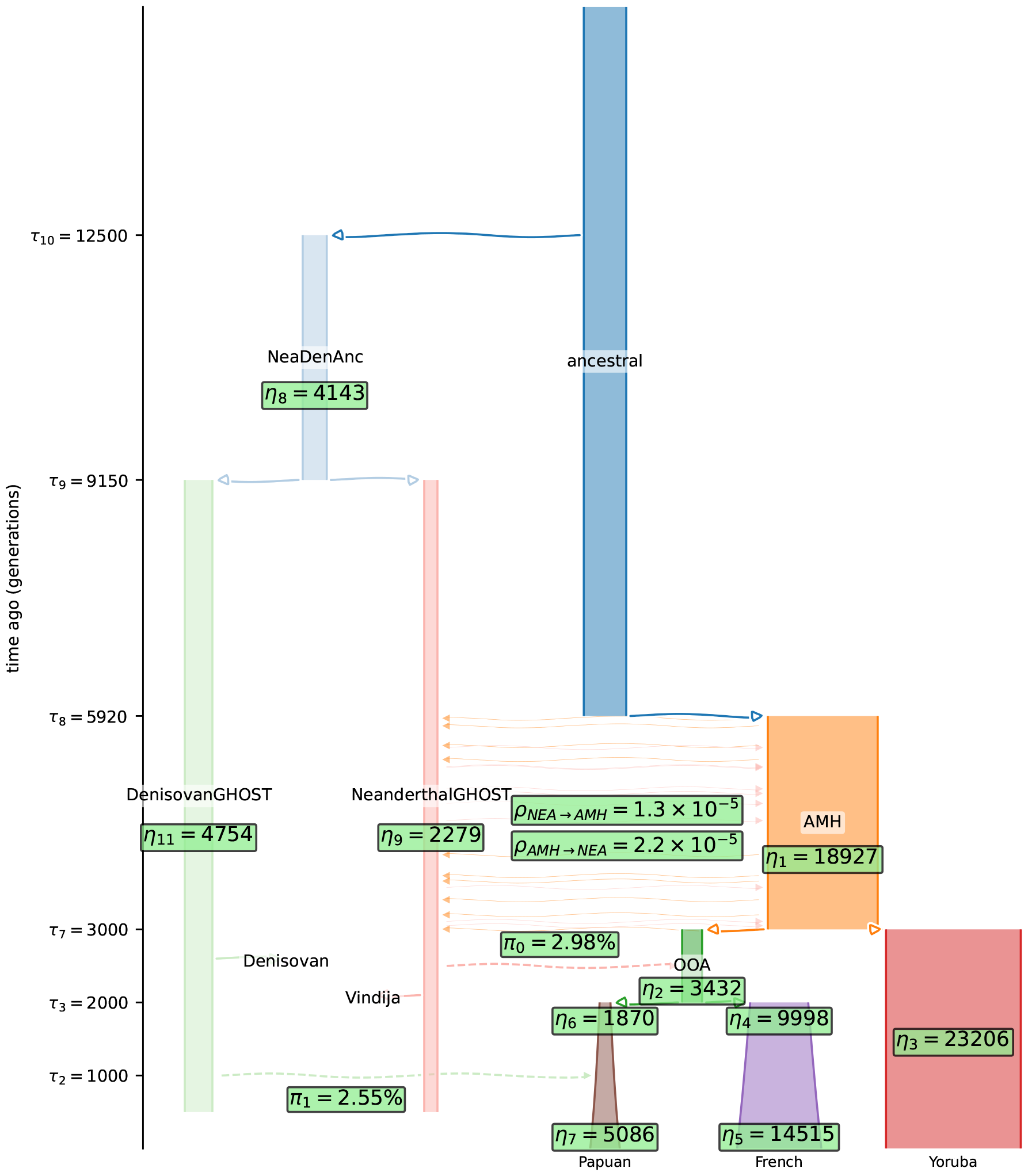
Out-of-Africa model with two extinct hominids and continuous gene flow between AMH and Neandertals. Parameter estimates are shown in green. *η* parameters are effective population sizes (2N), *π* parameters are admixture proportions and *ρ* parameters are migration rates.

There are a total of 12 parameters in the model, summarized in Table 4. (We did not attempt to estimate time parameters, and fixed them to the values originally inferred by Reich et al.) One hundred non-parametric bootstrap replicates, formed by sampling with replacement from the observed frequency spectrum, were also performed in order to obtain confidence intervals. Each run took about ten minutes using a Tesla V100 GPU.

**Table 4:** Estimated parameters along with their nonparametric bootstrap confidence intervals of the demography in Figure 3.

| Parameter | Description | Estimate | 2.5% | 97.5% |
| --- | --- | --- | --- | --- |
| $\eta_1$ | Size of Ancient Modern Humans | 18928 | 18732 | 19155 |
| $\eta_2$ | Size of Out-of-Africa Humans | 3433 | 3371 | 3477 |
| $\eta_3$ | Size of Yoruba | 23207 | 23040 | 23363 |
| $\eta_4$ | Size of French after Out-of-Africa | 9998 | 9263 | 10707 |
| $\eta_5$ | Recent size of French | 14516 | 13744 | 15450 |
| $\eta_6$ | Size of Papuan after Out-of-Africa | 1870 | 1788 | 1967 |
| $\eta_7$ | Recent size of Papuan | 5087 | 4745 | 5350 |
| $\eta_8$ | Size of Neanderthal-Denisovan Ancestor | 4143 | 4007 | 4262 |
| $\eta_9$ | Size of Neanderthal | 2280 | 2201 | 2372 |
| $\eta_{11}$ | Size of Denisovan | 4754 | 4370 | 5131 |
| $\pi_0$ | Neanderthal Admixture proportion of OOA | 298.05% | 288.06% | 305.79% |
| $\pi_1$ | Denisovan Admixture proportion of Papuan | 254.56% | 242.77% | 266.63% |

Results are shown in the right columns of Table 4, with point estimates superimposed on the demography visualization (Figure 3) for convenience. In (Reich et al., 2010), Vindija Neanderthal admixture is estimated between 1.3%-3.7% and Denisovan admixture between 3.8%-5.8%. Our confidence interval for the Neanderthal individual agrees with their *S*-statistic based confidence interval (π_0_ at Table 4), however we find a a lower admixture proportion from Denisovan to Papuans (π_1_ at Table 4). Our findings agree with another study by (Vernot, Tucci, et al., 2016), who estimated the proportion of Denisovan to Papuan admixture to be around 3% using *f*_4_ analysis.

Next, we considered an extended model that also allowed for continuous gene flow between Neanderthals and anatomically modern humans. At present, the timing and nature of ancient human migration events, particularly those involving Neanderthals and modern humans, remain uncertain. To study this further, we added two additional parameters to the model considered above, which allows for a constant rate of asymmetric migration between AMH and Neandertal between approximately 100,000 and 175,000 years ago.

Adding continuous migration to the model requires considerably more computation. The time needed to evaluate the gradient of the log-likelihood jumped from around one second in the pulse-only model, to ∼200s. This was too long for bootstrapping to be feasible, so instead we obtained maximum likelihood estimates by running a single optimization, and used asymptotic theory to form confidence intervals. Results are shown in Table 5. The migration rate from Ancient Modern Humans to Neanderthals (ρ_1_) was found to be almost twice the rate of the other direction (ρ_0_), indicating a greater gene flow from Ancient Modern Humans to Neanderthals. ρ_1_ also had a narrower confidence interval compared to ρ_0_.

**Table 5:** Estimated migration rates along with their parametric confidence intervals of the demography in Figure 4.

| Parameter | Description | Estimate | 2.5% | 97.5% |
| --- | --- | --- | --- | --- |
| $\rho_0$ | Migration rate from Neanderthal to AMH | 1.25e-05 | 1.04e-05 | 1.46e-05 |
| $\rho_1$ | Migration rate from AMH to Neanderthal | 2.24e-05 | 2.17e-05 | 2.31e-05 |

To better understand this asymmetry, we exploited momi3’s ability to quickly evaluate the log-likelihood function for many parameter values, and directly visualized the likelihood surface (Figure 5a). The log-likelihood values were calculated on a grid of migration rates around their maximum likelihood estimates. The values of log-likelihoods represent the values subtracted from the log-likelihood without migration (value for 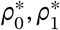 is equal to 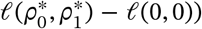). The green line represents the likelihood values while holding ρ_0_ constant at 2.24 × 10^−5^, and the red line represents the likelihood values while holding ρ_0_ constant at 1.25 × 10^−5^. We plotted these lines in Figure 5b. We observed that ρ_1_ had a sharper peak, resulting in a narrower confidence interval in Table 5. The flatter peak on ρ_0_ (Neanderthal to AMH) indicates that even small values of ρ_0_ are still relatively likely. Hence, the migration rate from Ancient Modern Humans to Neanderthals seems much more pronounced compared to the other direction.

**Figure 5.**
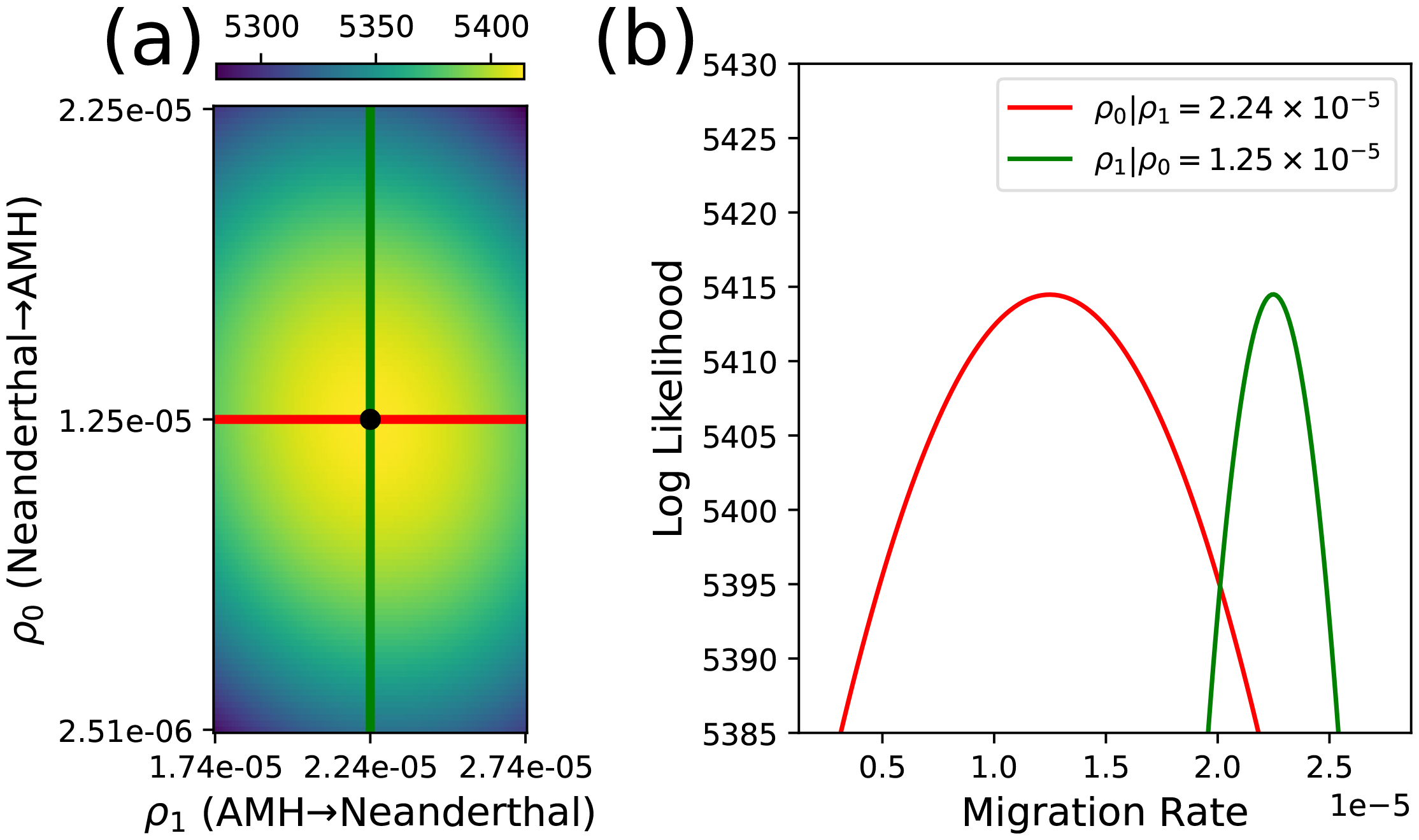
The likelihood surface of migration rates is depicted in (a), where the likelihood is evaluated using a grid of migration rates and visualized through a heatmap. The likelihood is represented by lighter colors indicating higher values. In (b), the migration rate is plotted against the likelihood for both migration parameters, while holding the other migration parameter constant at its maximum likelihood estimate (MLE).

## 4 Conclusion

We have described a new software package, momi3, for inferring complex demographies from the joint site frequency spectrum of multiple populations. This work extends our earlier method momi2 in several ways. First, we added support for continuous migration, bringing the feature set of momi at least to parity with other methods in this space. Second, we improved on the existing state-of-the-art by implementing a novel “genealogical importance sampling” strategy, which prunes unlikely genealogies from the analysis, reducing memory and computational requirements for large sample sizes. Finally, we made various implementation and user interface improvements which make momi3 easier to use and able to run seamlessly on CPUs and GPUs.

On simulated data, momi3 is up to 1000× faster than existing methods, with more modest speed improvements depending on the nature of the underlying demography. We also applied momi3 to real data, analyzing multiple genomes to study archaic admixture in modern humans. We constructed a demography similar the one studied by Reich et al. (2010) and fit it to all the available data in Wohns et al. (2022). To the best of our knowledge, no existing method could successfully fit such a model to a dataset of this size.

There are several areas for future improvement. Due to the currently experimental support for sparse tensor operations in Jax, momi3 is not that much faster, if at all, than existing methods (in particular moments) for analyzing demographies that feature continuous migration between a large number of populations. We expect that performance will improve this portion of the Jax code base matures. Similarly, JIT compilation times using our method can be long: comparing Table 2 with Table 1, we see that compilation times are currently approximately 20 times slower than the actual runtimes. While the one-time cost of compilation is generally negligible in optimization problems that require running the optimizer multiple times until convergence, it becomes a significant issue when experimenting with different demographies. In the case of momi3, recompilation is necessary every time the event tree changes. Thus, the lengthy compilation times become a hindrance when scientists aim to explore diverse population graphs. However, if future advancements in Jax reduce the compilation times, researchers would be able to quickly explore different population graphs, effectively expanding the space of analyzable models.

In the context of migration analysis, we have acknowledged and it has been previously noted by Gravel et al., that the moments quantity Φ_*n*_ bears a strong resemblance to the partial likelihoods tracked by momi, exhibiting a “dual” relationship. This observation suggests the potential for integrating these two methods into a unified approach, combining the advantages of both. For instance, such integration could facilitate the incorporation of selection and dominance effects within the framework of momi.

The work presented here focuses on studying genetically isolated subpopulations evolving on population genetic timescales. However, almost the same methodology could be applied to analyze phylogenetic data. Indeed, the computations needed to evaluate the likelihood of character data evolving along a phylogenetic reticulate network are formally identical to those used to study demographies featuring multiple pulse admixtures (Huson and Bryant, 2006; J. Zhu et al., 2018). To make momi3 useful for phylogenetic analysis, we need to allow for the possibility of recurrent mutations, and shift from a bi-to a tetra-allelic mutation model. This is technically feasible, but requires extensively modifying our algorithms and code, and is left to future work.

## Acknowledgements

The authors thank Jack Kamm for providing helpful feedback on a draft of the manuscript. This research was supported by NSF grant DMS-2052653 and the National Institute of General Medical Sciences of the NIH under award number R35GM151145. The content is solely the responsibility of the author and does not necessarily represent the official views of the NIH.

## Supplemental Tables

**Table S1:** Parameters in Constant Population Size Models.

| Parameters | 2D | 3D | 4D | 5D |
| --- | --- | --- | --- | --- |
| Size of ANC | ✓ | ✓ | ✓ | ✓ |
| Size of A | ✓ | ✓ | ✓ | ✓ |
| Size of B | ✓ | ✓ | ✓ | ✓ |
| Size of C |  | ✓ | ✓ | ✓ |
| Size of D |  |  | ✓ | ✓ |
| Size of E |  |  |  | ✓ |
| Split time | ✓ | ✓ | ✓ | ✓ |
| **Total number of parameters** | 4 | 5 | 6 | 7 |

**Table S2:** Parameters in Constant Population Size Models.

| **Parameters** | **OOA 2D**<br>**Migrations** | **OOA 3D**<br>**Migrations** | **OOA 4D**<br>**Migrations** | **OOA 5D**<br>**Migrations** | **OOA 5D**<br>**Pulses** |
| --- | --- | --- | --- | --- | --- |
| Ancient size of YRI | ✓ | ✓ | ✓ | ✓ | ✓ |
| Recent size of YRI | ✓ | ✓ | ✓ | ✓ | ✓ |
| Size of Neanderthal |  |  |  | ✓ | ✓ |
| Size of ArchaicAFR |  |  | ✓ | ✓ | ✓ |
| Ancient size of CEU | ✓ | ✓ | ✓ | ✓ | ✓ |
| Size of CEU just before exp growth | ✓ | ✓ | ✓ | ✓ | ✓ |
| Recent size of CEU | ✓ | ✓ | ✓ | ✓ | ✓ |
| Size of CHB just before exp growth |  | ✓ | ✓ | ✓ | ✓ |
| Recent size of CHB |  | ✓ | ✓ | ✓ | ✓ |
| Migration rate between YRI and ArchaicAFR |  |  | ✓ | ✓ |  |
| Migration rate (1st) between YRI and CEU | ✓ | ✓ | ✓ | ✓ |  |
| Migration rate (2nd) between YRI and CEU | ✓ | ✓ | ✓ | ✓ |  |
| Migration rate between CEU and Neanderthal |  |  |  | ✓ |  |
| Migration rate between CHB and CEU |  | ✓ | ✓ | ✓ |  |
| Migration rate between CHB and Neanderthal |  |  |  | ✓ |  |
| CHB-CEU split time & Recent migration start time | ✓ | ✓ | ✓ | ✓ | ✓ |
| Time of pulse migration from Neanderthal to CEU |  |  |  |  | ✓ |
| CEU-YRI split time | ✓ | ✓ | ✓ | ✓ | ✓ |
| Time of pulse migration from YRI to Neanderthal |  |  |  |  | ✓ |
| Start time of YRI-ArchaicAFR migration |  |  | ✓ | ✓ |  |
| Size change time of YRI | ✓ | ✓ | ✓ | ✓ | ✓ |
| ArchaicAFR-YRI split time |  |  | ✓ | ✓ | ✓ |
| Neanderthal-YRI split time |  |  |  | ✓ | ✓ |
| Pulse migration from Neanderthal to CEU |  |  |  |  | ✓ |
| Pulse migration from YRI to Neanderthal |  |  |  |  | ✓ |
| **Total number of parameters** | 10 | 13 | 17 | 21 | 18 |

## Supplemental Figures

**Figure S1:**
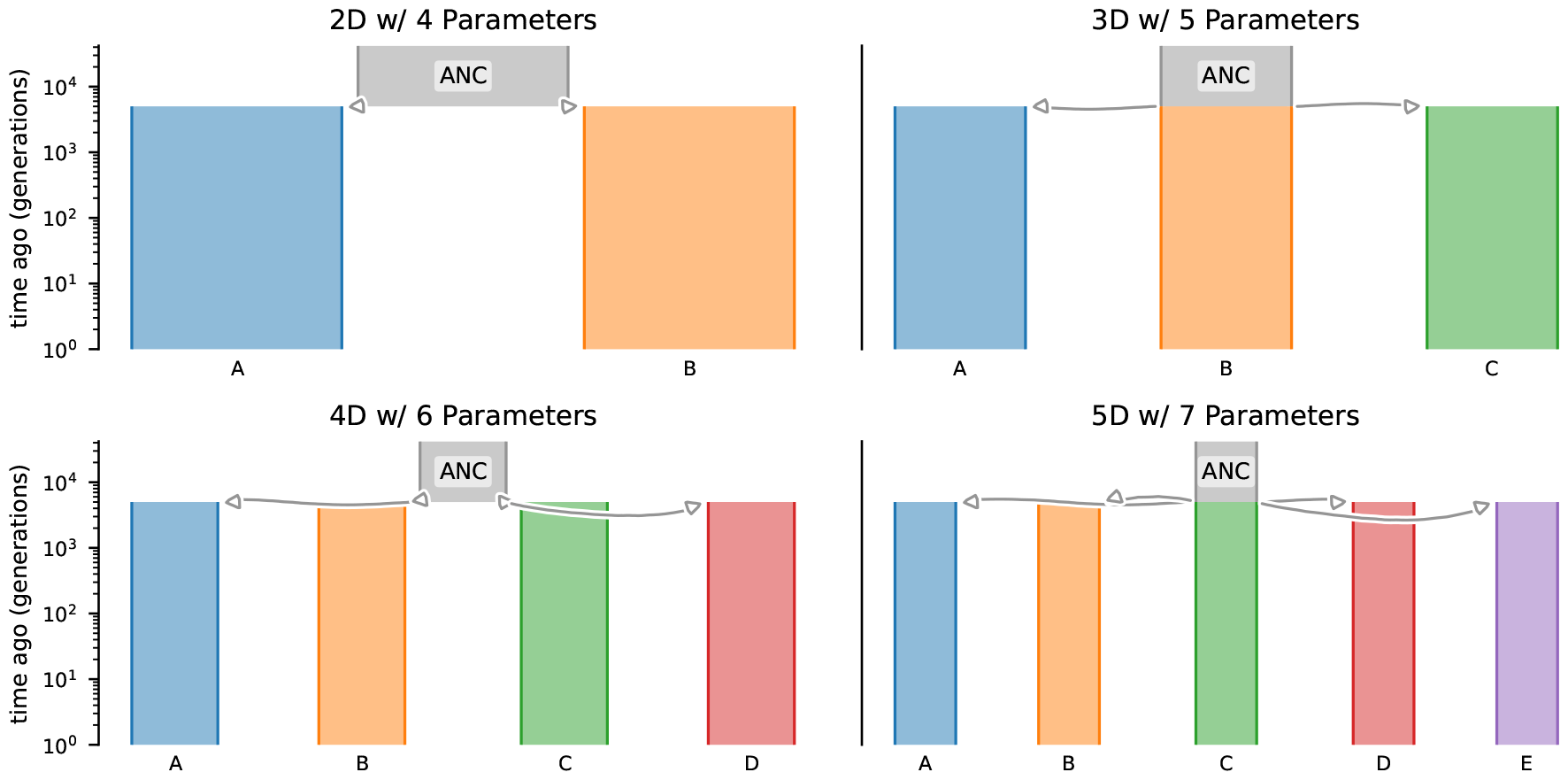
Constant population size demographies. All demes have population size of 2*N* = 5 × 10^3^, and they diverged 5 × 10^3^ generations ago. Parameters can be seen at Table S1.

**Figure S2:**
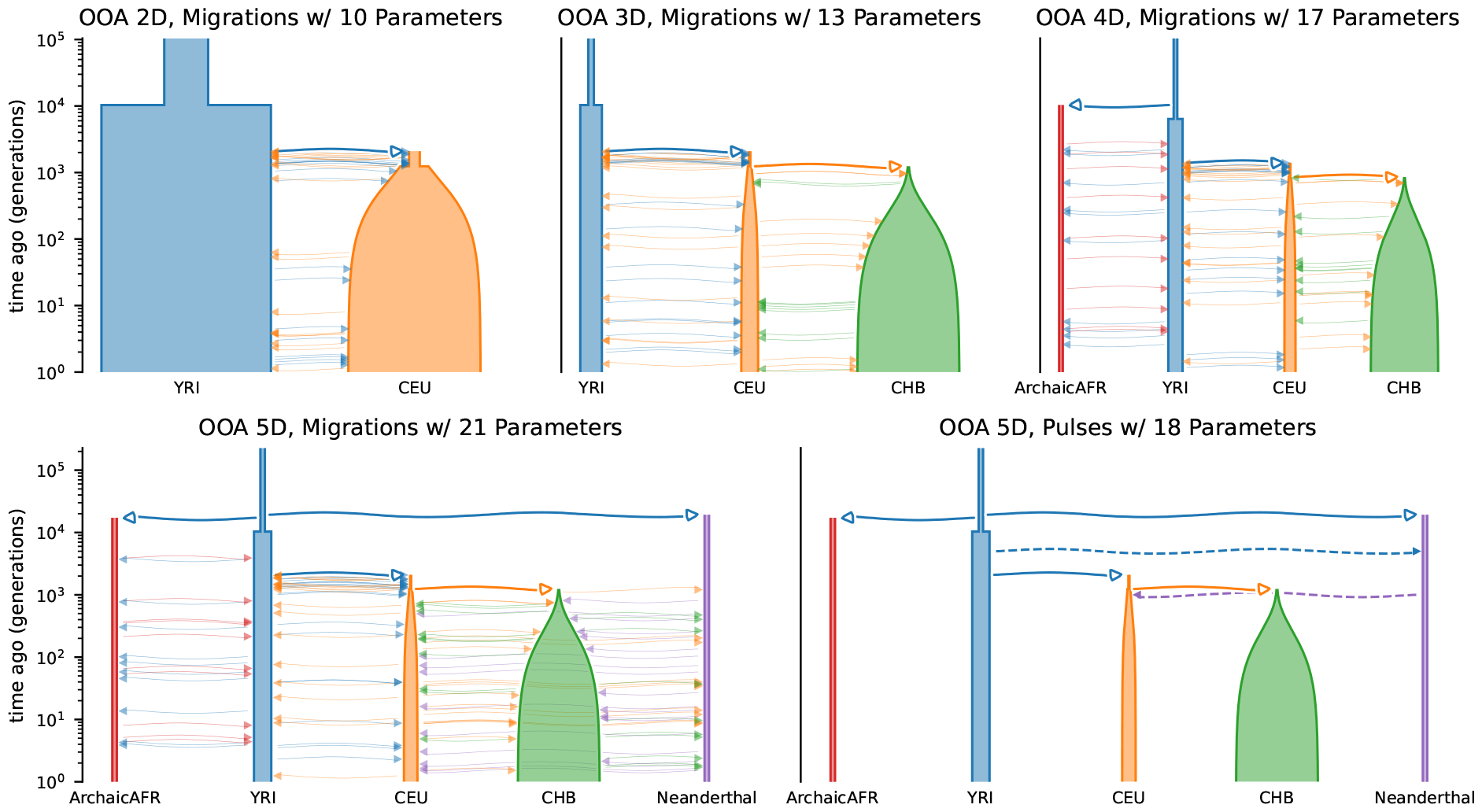
Out-of-Africa demographies. Parameters can be seen at Table S2.

**Figure S3:**
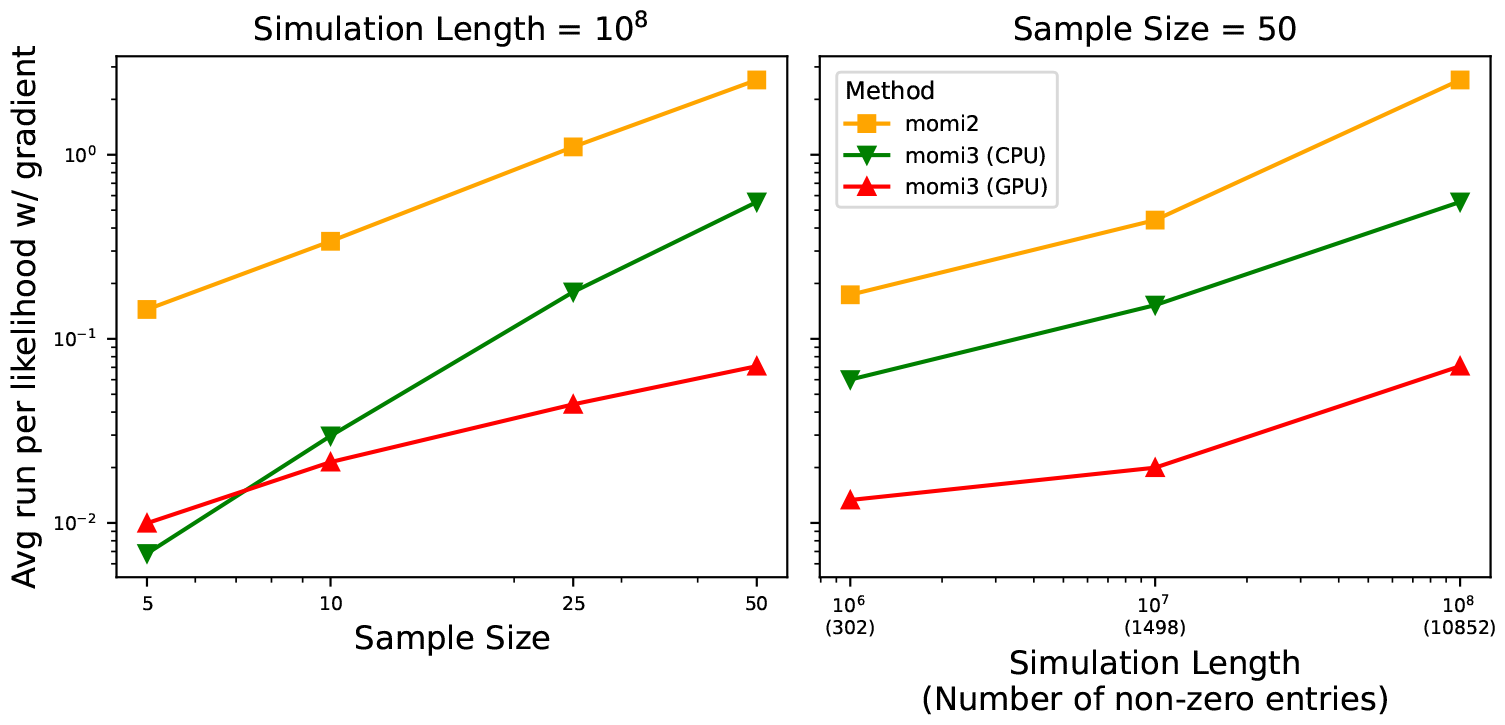
Runtime of a likelihood with gradient for the demography *4D w/ 6 Parameters* (See Figure S1). The figure shows how each method scales by the sample size (left) and simulation length (right).

**Figure S4:**
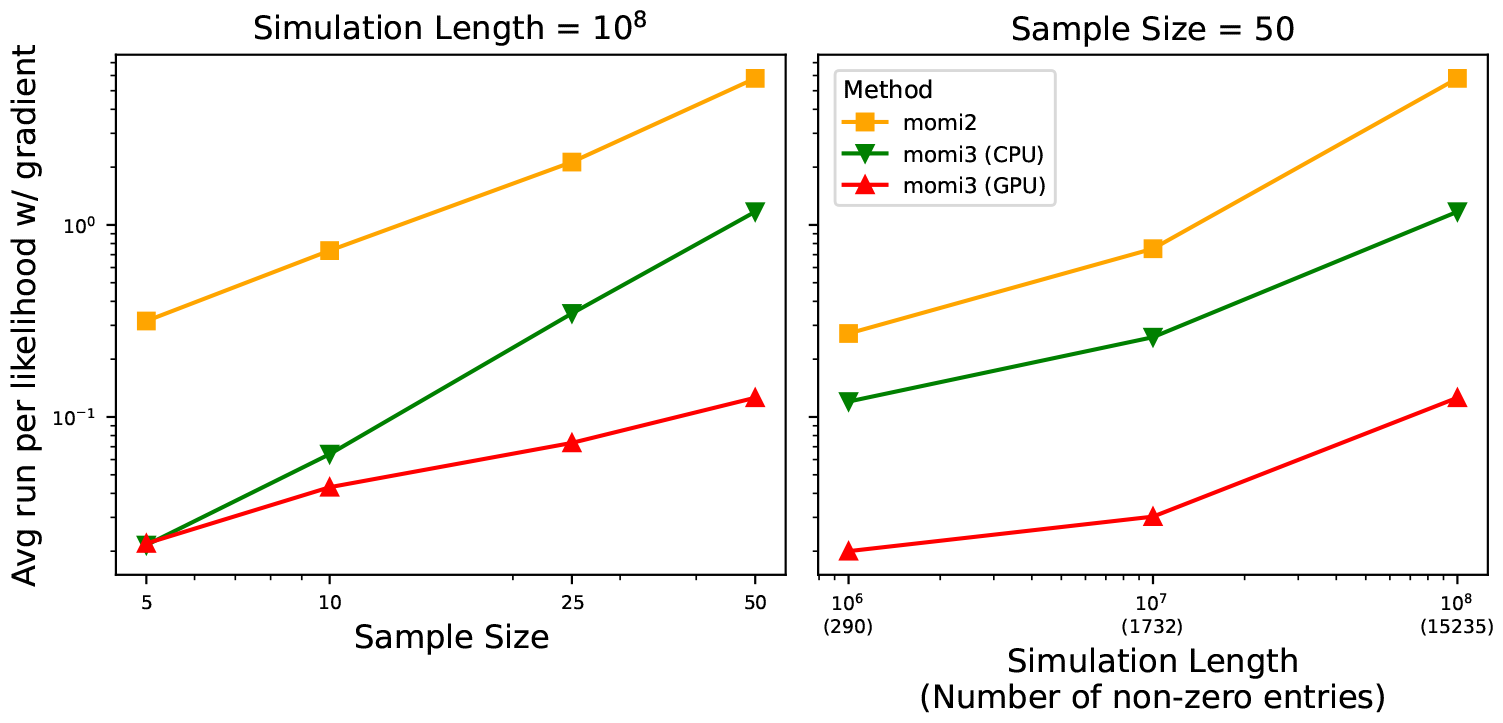
Runtime of a likelihood with gradient for the demography *5D w/ 7 Parameters* (See Figure S1). The figure shows how each method scales by the sample size (left) and sequence length (right).

**Figure S5:**
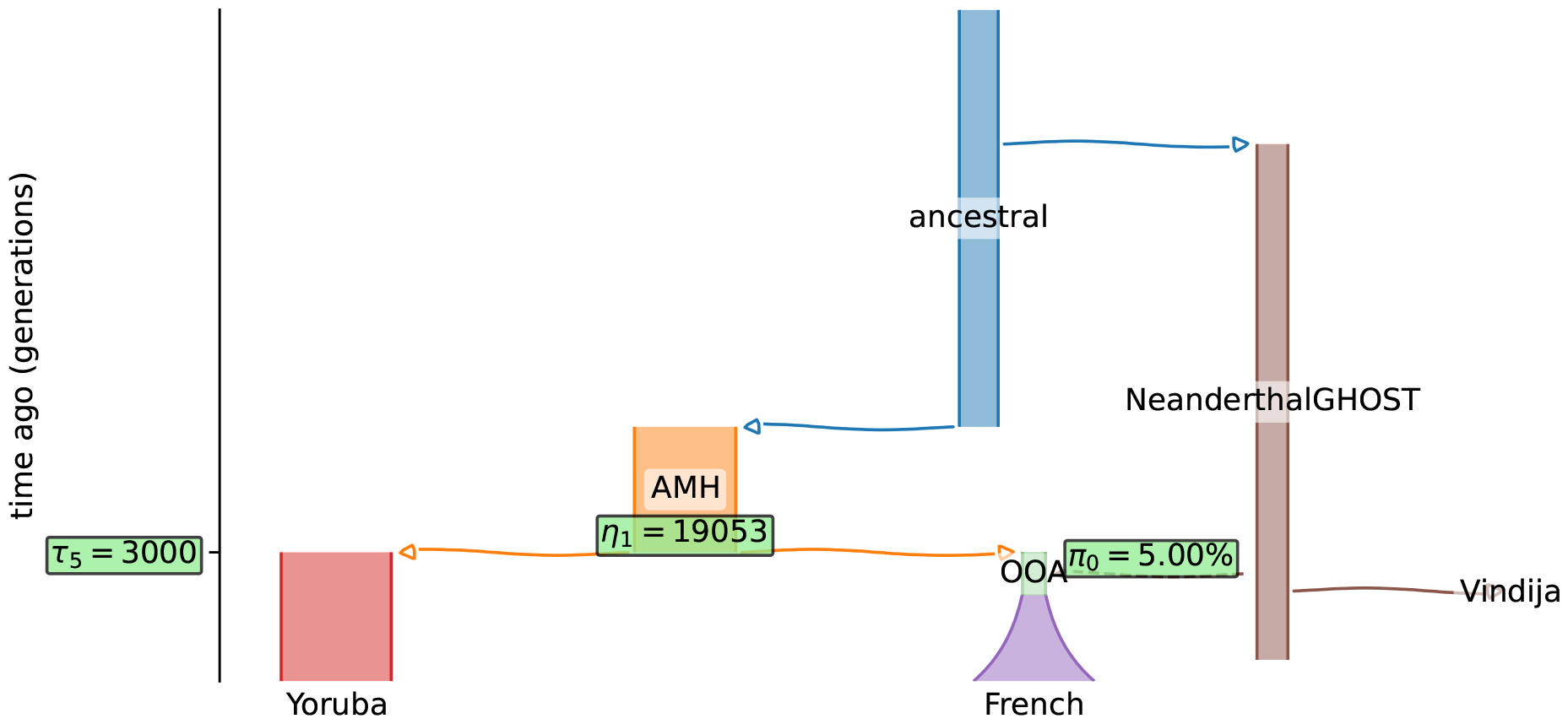
Out-of-Africa demography with Neanderthal admixture. Variables highlighted in green are trained, while the remaining parameters of the demography were fixed. The trained variables wee: *τ*_5_ is the time of Out-of-Africa (OOA) event; *η*_1_ is the population size of Ancient Modern Humans; *π*_0_ is the admixture fraction of Neanderthal in the OOA population.

**Figure S6:**
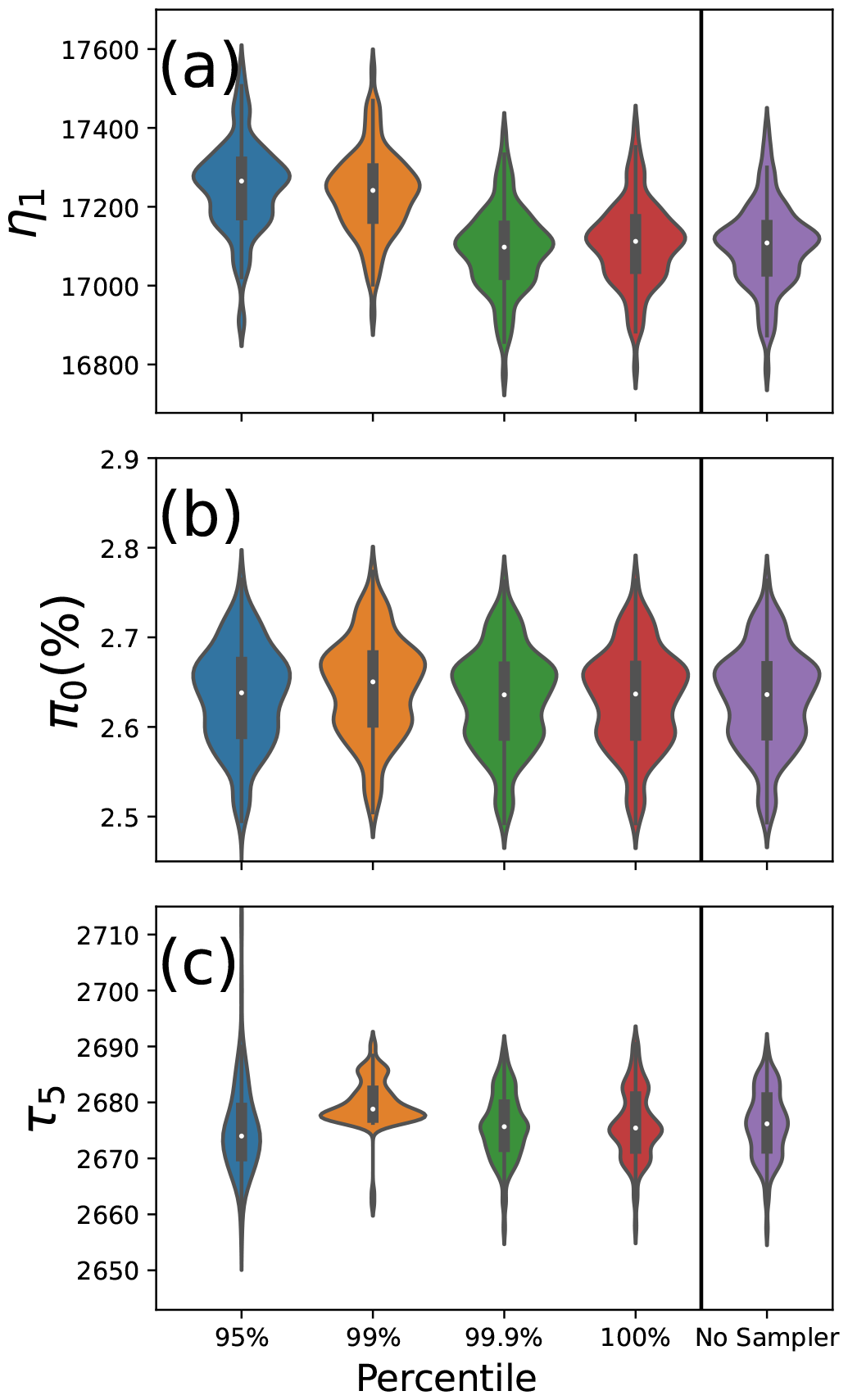
Distribution of the estimates of Figure 4 model in violin plots. Distributions are obtained by bootstrap samples. Each percentiles are used by the genealogical importance sampler. We compared them with the No Sampler. Closer to No Sampler is better.

## Footnotes

1 Instead of simulating, we could also employ exact expressions for the distribution of 𝒜 _*n*_(*t*) (S. Tavaré, 1984), however these are known to be numerically unstable for large *n*.

2 Note that, due to Monte Carlo error, setting ϵ = 0 (i.e., the max over all Monte Carlo trials) still results in performance improvements, and is not equivalent to the “No Sampling” case.

## References

Bartlett, Roscoe (2008). A derivation of forward and adjoint sensitivities for ODEs and DAEs. Tech. rep. Tech. rep., Tech. Rep. SAND2007-6699, Sandia National Laboratories.

Baumdicker, Franz et al. (2022). “Efficient ancestry and mutation simulation with msprime 1.0”. In: Genetics 220.3, iyab229.

Bhaskar, A., Y. X. Rachel Wang, and Y. S. Song (2015). “Efficient inference of population size histories and locus-specific mutation rates from large-sample genomic variation data”. In: Genome Research 25.2, pp. 268–279.

Bradbury, James et al. (2018). JAX: composable transformations of Python+NumPy programs. Version 0.2.5. url: http://github.com/google/jax.

Browning, Sharon R et al. (2018). “Analysis of human sequence data reveals two pulses of archaic Denisovan admixture”. In: Cell 173.1, pp. 53–61.

Chen, Hua (2012). “The joint allele frequency spectrum of multiple populations: A coalescent theory approach”. In: Theoretical Population Biology 81.2, pp. 179– 195.

Chen, Hua (2013). “Intercoalescence Time Distribution of Incomplete Gene Genealogies in Temporally Varying Populations, and Applications in Population Genetic Inference”. In: Annals of Human Genetics 77.2, pp. 158–173.

Dannemann, Michael and Fernando Racimo (2018). “Something old, something borrowed: admixture and adaptation in human evolution”. In: Current opinion in genetics & development 53, pp. 1–8.

Durvasula, Arun and Sriram Sankararaman (2020). “Recovering signals of ghost archaic introgression in African populations”. In: Science Advances 6.7, eaax5097.

Excoffier, Laurent, Isabelle Dupanloup, et al. (2013). “Robust Demographic Inference from Genomic and SNP Data”. In: PLoS Genetics 9.10, e1003905.

Excoffier, Laurent and Matthieu Foll (2011). “Fastsimcoal: a continuous-time coalescent simulator of genomic diversity under arbitrarily complex evolutionary scenarios”. In: Bioinformatics 27.9, pp. 1332–1334.

Excoffier, Laurent, Nina Marchi, et al. (2021). “fastsimcoal2: demographic inference under complex evolutionary scenarios”. In: Bioinformatics 37.24, pp. 4882– 4885.

Gower, Graham et al. (2022). “Demes: a standard format for demographic models”. In: Genetics 222.3, iyac131.

Green, Richard E et al. (2010). “A draft sequence of the Neandertal genome”. In: Science 328.5979, pp. 710–722.

Griffiths, R.C. and Simon Tavaré (1998). “The age of a mutation in a general coalescent tree”. In: Communications in Statistics. Stochastic Models 14.1-2, pp. 273–295.

Gutenkunst, Ryan N. (2021). “dadi. CUDA: Accelerating population genetics inference with graphics processing units”. In: Molecular biology and evolution 38.5, pp. 2177–2178.

Gutenkunst, Ryan N. et al. (2009). “Inferring the Joint Demographic History of Multiple Populations from Multidimensional SNP Frequency Data”. In: PLoS Genetics 5.10, e1000695.

Huson, Daniel H and David Bryant (2006). “Application of phylogenetic networks in evolutionary studies”. In: Molecular biology and evolution 23.2, pp. 254–267.

Jouganous, Julien et al. (July 2017). “Inferring the Joint Demographic History of Multiple Populations: Beyond the Diffusion Approximation”. en. In: Genetics 206.3, pp. 1549–1567. ISSN: 0016-6731, 1943-2631. doi: 10.1534/genetics.117.200493.

Kamm, John A, Jonathan Terhorst, Richard Durbin, et al. (2020). “Efficiently inferring the demographic history of many populations with allele count data”. en. In: J. Am. Stat. Assoc. 115.531, pp. 1472–1487. ISSN: 0162-1459. doi: 10.1080/01621459.2019.1635482.

Kamm, John A, Jonathan Terhorst, and Yun S Song (Feb. 2017). “Efficient computation of the joint sample frequency spectra for multiple populations”. en. In: J. Comput. Graph. Stat. 26.1, pp. 182–194. ISSN: 1061-8600. doi: 10.1080/10618600.2016.1159212.

Kelleher, Jerome, Alison M Etheridge, and Gilean McVean (2016). “Efficient coalescent simulation and genealogical analysis for large sample sizes”. In: PLoS computational biology 12.5, e1004842.

Li, N. and M. Stephens (2003). “Modeling Linkage Disequilibrium and Identifying Recombination Hotspots Using Single-Nucleotide Polymorphism Data”. In: Genetics 165, pp. 2213–2233.

Mallick, Swapan et al. (2016). “The Simons genome diversity project: 300 genomes from 142 diverse populations”. In: Nature 538.7624, pp. 201–206.

Marchi, Nina, Flávia Schlichta, and Laurent Excoffier (2021). “Demographic inference”. In: Current Biology 31.6, R276–R279.

Meyer, Matthias et al. (2012). “A high-coverage genome sequence from an archaic Denisovan individual”. In: Science 338.6104, pp. 222–226.

Al-Mohy, Awad H. and Nicholas J. Higham (2011). “Computing the Action of the Matrix Exponential, with an Application to Exponential Integrators”. In: SIAM Journal on Scientific Computing 33 (2), pp. 488–511.

Prüfer, Kay et al. (2014). “The complete genome sequence of a Neanderthal from the Altai Mountains”. In: Nature 505.7481, pp. 43–49.

Ragsdale, Aaron P and Simon Gravel (2019). “Models of archaic admixture and recent history from two-locus statistics”. In: PLoS genetics 15.6, e1008204.

Rasmussen, Matthew D et al. (2014). “Genome-wide inference of ancestral recombination graphs”. In: PLoS Genetics 10.5, e1004342.

Reich, David et al. (2010). “Genetic history of an archaic hominin group from Denisova Cave in Siberia.” In: Nature 468.7327, pp. 1053–1060.

Sankararaman, Sriram et al. (2014). “The genomic landscape of Neanderthal ancestry in present-day humans”. In: Nature 507.7492, pp. 354–357.

Sheehan, Sara, Kelley Harris, and Yun S Song (2013). “Estimating variable effective population sizes from multiple genomes: A sequentially Markov conditional sampling distribution approach”. In: Genetics 194.3, pp. 647–662.

Tavaré, S. (1984). “Line-of-descent and genealogical processes, and their applications in population genetics models”. In: Theor. Popul. Biol. 26, pp. 119–164.

Terhorst, Jonathan, John A Kamm, and Yun S Song (2017). “Robust and scalable inference of population history from hundreds of unphased whole genomes”. In: Nature genetics 49.2, pp. 303–309.

The 1000 Genomes Project Consortium (2015). “A global reference for human genetic variation”. In: Nature 526.7571, pp. 68–74.

Vernot, Benjamin and Joshua M Akey (2015). “Complex history of admixture between modern humans and Neandertals”. In: The American Journal of Human Genetics 96.3, pp. 448–453.

Vernot, Benjamin, Serena Tucci, et al. (2016). “Excavating Neandertal and Denisovan DNA from the genomes of Melanesian individuals”. In: Science 352.6282, pp. 235–239. doi: 10.1126/science.aad9416.

Villanea, Fernando A and Joshua G Schraiber (2019). “Multiple episodes of interbreeding between Neanderthal and modern humans”. In: Nature ecology & evolution 3.1, pp. 39–44.

Wainwright, Martin J and Michael Irwin Jordan (2008). Graphical models, exponential families, and variational inference. Now Publishers Inc.

Wohns, Anthony Wilder et al. (2022). “A unified genealogy of modern and ancient genomes”. In: Science 375.6583, eabi8264.

Zhu, Jiafan et al. (2018). “Bayesian inference of phylogenetic networks from biallelic genetic markers”. In: PLoS computational biology 14.1, e1005932.

